# Pathogenic CANVAS (AAGGG)_n_ repeats stall DNA replication due to the formation of alternative DNA structures

**DOI:** 10.1101/2023.07.25.550509

**Authors:** Julia A. Hisey, Elina A. Radchenko, Silvia Ceschi, Anastasia Rastokina, Nicholas H. Mandel, Ryan J. McGinty, Gabriel Matos-Rodrigues, Alfredo Hernandez, André Nussenzweig, Sergei M. Mirkin

## Abstract

CANVAS is a recently characterized repeat expansion disease, most commonly caused by homozygous expansions of an intronic (A_2_G_3_)_n_ repeat in the *RFC1* gene. There are a multitude of repeat motifs found in the human population at this locus, some of which are pathogenic and others benign. In this study, we conducted structure-functional analyses of the main pathogenic (A_2_G_3_)_n_ and the main nonpathogenic (A_4_G)_n_ repeats. We found that the pathogenic, but not the nonpathogenic, repeat presents a potent, orientation-dependent impediment to DNA polymerization *in vitro*. The pattern of the polymerization blockage is consistent with triplex or quadruplex formation in the presence of magnesium or potassium ions, respectively. Chemical probing of both repeats in supercoiled DNA reveals triplex H-DNA formation by the pathogenic repeat. Consistently, bioinformatic analysis of the S1-END-seq data from human cell lines shows preferential H-DNA formation genome-wide by (A_2_G_3_)_n_ motifs over (A_4_G)_n_ motifs *in vivo*. Finally, the pathogenic, but not the non-pathogenic, repeat stalls replication fork progression in yeast and human cells. We hypothesize that CANVAS-causing (A_2_G_3_)_n_ repeat represents a challenge to genome stability by folding into alternative DNA structures that stall DNA replication.

## INTRODUCTION

Biallelic expansions of (A_2_G_3_)_n_ repeats cause a newly discovered repeat expansion disease (RED) named CANVAS (**c**erebellar **a**taxia, **n**europathy, **v**estibular **a**reflexia **s**yndrome) ^1, 2^. It is an autosomal recessive disease with a carrier frequency range from 0.7% to 4% in studied populations, resulting in a prevalence range from 1:20,000 to 1:625, respectively ^1, 3^. This frequency establishes *RFC1*-related ataxia as likely the most common cause of hereditary late-onset ataxia ^1, 2, 4^. CANVAS is a progressive neurodegenerative disease with a mean age of onset of 52 and a broad spectrum of clinical features, including but not limited to: imbalance, peripheral sensory symptoms, oscillopsia, dry cough, autonomic dysfunction, dysarthria, and dysphagia ^4, 5^.

The expandable (A_2_G_3_)_n_ repeat resides in the poly-A tail of an AluSx3 element within the second intron of the *RFC1* gene ^1^. *RFC1* encodes the largest subunit of replication factor C, the complex responsible for loading PCNA onto DNA during DNA replication and repair ^6, 7^. Given its essential role, it is not surprising that this is the first *RFC1* mutation found to cause human disease. While the pathogenic mechanism of CANVAS remains uncertain, RFC1 loss of function is suspected due to (1) its recessive inheritance and (2) the discovery of CANVAS-affected patients heterozygous for the repeat expansion and an *RFC1* truncating mutation ^3, 8–11^. CANVAS’s disease pathogenesis is an area of intense biomedical research and at the heart of every RED’s pathogenesis and genetics are the repeats themselves.

Most expandable repeats can adopt alternative, non-B DNA secondary structures that are integral to their propensity to expand and cause disease (reviewed in ^12^). These secondary structures are formed during processes involving B-DNA unwinding, such as replication, transcription, and repair, and, at the same time, they can become an obstacle for these processes. Notably, DNA replication and repair have been identified as major sources of repeat instability across various repeats, which has been attributed to their structure-prone nature (reviewed in ^12^).

CANVAS differs from most other REDs in that its pathogenic allele varies from the nonpathogenic one not only in repeat size, but also in its base composition. While most healthy individuals harbor (A_4_G)_11-100_ repeats in the *RFC1* locus, CANVAS patients largely carry (A_2_G_3_)_250-2000_ repeats ^1, 13^. Other iterations of the repeat exist, and their pathogenicity is currently being unraveled. Oftentimes, repeat variants are not pure and contain expanded (A_2_G_3_)_n_ or other repeat interruptions, adding complexity to assigning the alternative repeat to the pathogenic or benign category ^13^. Alternative repeat motifs include (A_3_G_2_)_n_, (ACAGG)_n_, (AGGGC)_n_, (AAGGC)_n_, (AAGAG)_n_, (AGAGG)_n_, (AAAGGG)_n_, (ACGGG)_n_, (AG_4_)_n_, and (AACGG)_n_ ^1, 13–18^. Interestingly, the propensity of the most frequently found repeats to expand correlates with the number of guanine residues in the repetitive unit: A_4_G < A_3_G_2_< A_2_G_3_ ^1^. Given the repeat composition, we hypothesized the pathogenic repeats can form a stable triplex H-DNA or G-quadruplex DNA, since they are simultaneously homopurine/homopyrimidine (hPu/hPy) mirror repeats ^19^ and contain evenly spaced G3 runs_20._

Thus, we set out to determine if the pathogenic (A_2_G_3_)_n_ repeats form an alternative DNA secondary structure(s), what this structure(s) is, and whether it impedes replication, which *a priori* may lead to the repeat’s instability. We found that pathogenic (A_2_G_3_)_10_ repeats, but not the nonpathogenic (A_4_G)_10_ repeats, strongly stall DNA Pol I polymerization and bacteriophage T7 polymerization *in vitro.* The stalling is orientation-dependent, occurring only when (A_2_G_3_)_n_ serves as the template strand. Ambient conditions have a profound effect on the position of the stall, with patterns suggestive of G-quadruplex formation in the presence of potassium or triplex formation in the presence of magnesium. Using chemical probing, we identified the formation of H-r triplex DNA (pyrimidine-purine-purine triplex)^19^ by the pathogenic (A_2_G_3_)_n_ repeat *in vitro*. Analysis of S1-END-seq peaks at genome-wide H-motifs in human cells revealed that (A_2_G_3_)_n_ tracts have a higher propensity than (A_4_G)_n_ tracts to form triplexes *in vivo*. Using two-dimensional electrophoretic analysis of replication intermediates, we show that the pathogenic repeat causes orientation-dependent replication stalling in both yeast and human cells when the (A_2_G_3_)_n_ run is in the lagging strand template. We suggest, therefore, that non-B DNA-forming potential of the pathogenic repeat during DNA replication results in fork stalling, which may have a role in its instability.

## MATERIALS AND METHODS

Details of construction of plasmids and strains used in this study are described in the Supplemental Materials. Specific PCR programs, plasmids, primers, and strains used in this study are listed in Supplemental Tables 1-6.

### Cloning and amplification of (A_2_G_3_)_n_ repeats

Due to the secondary structure-forming potential of these repeats, adjustments were made to PCR reaction mixes and cycling programs: Thermo Scientific Phusion High Fidelity DNA Polymerase (Cat# F530L) was used to amplify fragments for Gibson Assembly, cloning, or fragments for yeast transformation. The manufacturer’s reaction for Phusion PCR using was altered as follows: the master mix included 1M Betaine and no DMSO. The program listed in Supplemental Table 1 was followed with the following general changes: the extension time was lengthened to allow for progression through the entire repeat sequence and the extension temperature was raised to 75°C or 80°C when necessary. For some PCRs, the annealing temperature was also raised, and longer primers were designed accordingly to ensure annealing. For products with multiple bands used for cloning, the properly sized fragment was gel extracted and repeat length was confirmed with the Taq repeat PCR programs before cloning. Specific protocols are described in Supplemental Table 2 and generally, annealing and extension temperatures were raised and extension time lengthened. Phusion was also used to amplify the (A_4_G)_n_ repeats. Genescript Taq polymerase (Cat# E00007) was used to amplify the (A_2_G_3_)_n_ repeats. The following PCR reaction was used: 1X Taq Buffer with (NH_4_)_2_SO_4_ (Thermo Scientific Cat# B33), 1mM primers, 100mM dNTP, 1mM MgCl_2_, 1M betaine, 0.625 units Taq polymerase, and 1ng DNA. The PCR programs used for different repeats are listed in Supplemental Table 3.

Supercoiled plasmids used for two-dimensional electrophoresis or chemical probing were run alongside the linearized plasmid on a 0.8% agarose gel with 0.5µg/mL EtBr (conditions for 7.8kb plasmid) or 0.6% agarose gel with 1µg/mL EtBr (conditions for 12.4kb plasmid) to confirm that their monomeric state. *E. coli* strains with (A_2_G_3_)_n_-bearing plasmids were grown at 23°C to reduce instability during plasmid replication.

### DNA polymerization

Double-stranded plasmids pJH2 and pJH9 (ordered from Genescript) were used as DNA templates for *in vitro* polymerization experiments with (A_2_G_3_)_10_ or (A_4_G)_10_ repeats, respectively. Primers JH271 and JH272 were used to polymerize through the repeats with either the pyrimidine or purine repeat in the template strand, respectively. Reactions using ThermoSequenase were carried out as follows: The USBio Thermo Sequenase Cycle Sequencing Kit (Cat# 78500) was used according to the manufacturer’s 3’-dNTP internal label cycling sequencing instructions with the following alterations and specifications: Instead of conducting multiple rounds of labeling and extending, 5µg of each plasmid and 0.5pmol of primer were used to conduct only one cycle. The primer was pre-annealed to the plasmid in 11µl water at 95°C for 2 minutes and immediately submerged in an ice-water bath, then the labeling components were added and the primer was labeled at 60°C for 30 seconds with 0.5µl of 1.25mM [α-^32^P]dATP. Upon aliquoting this reaction into the four pre-aliquoted termination mixes, termination was carried out at 72°C for 5 minutes and 4µl of stop solution was added to each reaction, mixed, and incubated at 95°C for 5 minutes and immediately submerged into an ice-water bath. 8µl of each reaction was loaded onto a 6% polyacrylamide gel with 7.5M urea prepared according to manufacturer’s instructions (National Diagnostics SEQUAGEL SEQUENCING SYSTEM 2.2 (Cat# EC-833)). Reactions using Vent polymerase were carried out as follows: Vent (exo-) DNA polymerase (New England Biolabs Cat# M0257S) was used according to primer extension experiments described in ^21^ with the following exceptions: 0.5pmol of primer and 5µg of DNA were pre-annealed as described above, the ddNTP:dNTP ratios and concentrations used are described in ^22^, labeling was carried out at 60°C for 30 seconds, termination was carried out at 90°C for 10 minutes, and steps after termination are identical to those with ThermoSequenase.

Primer/templates used for *in vitro* extension by T7 DNA polymerase were formed by pre-annealing 2.5µM 5’-^32^P labeled primer JH270 with 3.125µM single-stranded oligonucleotides bearing either (A_2_G_3_)_10_ (JH267), (T_2_C_3_)_10_ (JH268), or (A_4_G)_10_ (JH269) repeats in 10 mM Tris-HCl, pH 7.5, 0.1 mM EDTA alone or with the addition of 50mM K^+^ and 10mM Mg^2+^, or 50mM NaCl, KCl, or LiCl by incubation at 95°C for 5 minutes, followed by gradual return to room temperature. Primer extensions reactions consisted of 10nM primer/template and 1 µM T7 DNA polymerase in 40 mM Tris-HCl, pH 7.5, 5mM dithiothreitol (DTT), and 55mM of the indicated monovalent metal chloride salt (Li^+^, K^+^, or Na^+^). DNA polymerization was initiated by addition of 10mM MgCl_2_, followed by incubation at room temperature for 1 minute, and reactions were quenched with formamide/EDTA loading dye. Primer extension products were resolved by electrophoresis on a denaturing 10% polyacrylamide gel.

### Chemical probing

2µg of the repeat-containing pJH2 or pJH9 plasmid was incubated in a 10mM Tris-HCl pH 7.5 2mM MgCl2 buffer with either 6mM KMnO4 or the same volume of water for 2 minutes at 37°C. The reaction was quenched with 1M β-mercaptoethanol, precipitated with ethanol, rinsed with 70% ethanol, and resuspended in 5µl water. The primer extension was carried out using the same protocol as was described in the *in vitro* polymerization methods with the following changes: the chemically probed DNA was pre-annealed with 0.5pmol of primer JH271 in a final volume of 5.5µl, the labeling reaction’s final volume was 8.75µl with the same ratios as described in the manufacturer’s instructions, and dNTPs were added to a final concentration of 75mM for each dNTP for extension.

### Analysis of S1-END-seq data

S1-END-seq data from five cell lines were obtained from a previous study ^23^, provided as BED files, which represented the output of the “Peak calling” stage. Custom Python scripts were written to perform the following analysis. Overlapping genomic coordinates from the five cell lines were combined into non-overlapping peaks. Coordinates were converted to GRCh38 using Liftover. A set of control coordinates were randomly generated, matching the S1-END-seq peaks in total number, the length of each peak, and the proportion located on each chromosome. S1-END-seq peaks and control coordinates were compared to a database of repetitive sequences generated previously ^24^. For each category or subtype of repeat, the proportion of those falling within 100 nucleotides of the S1-END-seq peaks or control coordinates was calculated for each repeat length. Comparison of distributions along repeat lengths were made by Wilcoxon signed-rank test, restricted to length bins containing at least 3 repeats. Linear best fit lines for each distribution were weighted to the total number of repeats in each length bin and were also restricted to bins containing at least 3 repeats.

### Analysis of replication intermediates by two-dimensional (2D) gel electrophoresis in yeast and human cells

Yeast cells: The plasmids used for yeast two-dimensional gel electrophoresis contain the repeats, a yeast 2µ origin of replication, and ampicillin resistance. Plasmids were named pJH1, pJH26, and pJH4 and have (A_2_G_3_)_60_, (C_3_T_2_)_60_, or (A_4_G)_60_ in the lagging strand template in relation to the yeast 2µ origin. Plasmids were transformed into yeast strain JAH231 and yeast replication intermediates were extracted, restriction digested, run on 2D gel, and analyzed all according to ^25^. Details of plasmid construction are in the supplemental materials.

Human cells: The plasmids used for human cell two-dimensional gel electrophoresis contain the repeats, the SV40 origin of replication, the gene encoding the large T antigen, and ampicillin resistance. Plasmids were named pJH5, pJH6, and pJH11 and have (A_2_G_3_)_60_, (C_3_T_2_)_60_, or (A_4_G)_60_ in the lagging strand template in relation to the SV40 origin of replication. Following the methods outlined in ^26^, plasmids were transfected into HEK293T cells, incubated for 48 hours, replication intermediates were extracted, and intermediates were run on a 2D gel. Details of plasmid construction are in the Supplemental Materials.

## RESULTS

### Pathogenic (A_2_G_3_)_n_ repeats strongly stall DNA polymerization *in vitro* in an orientation-dependent manner

Structure-forming repeats pose a significant obstacle to polymerases for DNA synthesis during DNA replication and repair ^27–33^. This phenomenon is believed to be a driver of repeat instability (reviewed in ^12, 34, 35^). During replication, the Okazaki initiation zone remains single-stranded, allowing for structure formation preferentially on the lagging, rather than leading, strand template. Accordingly, replication issues are more severe for many repeats when their structure-prone strand is on the lagging strand template, and they are particularly unstable in this orientation (reviewed in ^12^). To the best of our knowledge, no experimental data are available on the replication nor instability of the expandable CANVAS (A_2_G_3_)_n_ repeat.

To investigate polymerization through the repeats in either orientation *in vitro*, we used ThermoSequenase, a mutated DNA Polymerase I (exo-)^36^, and a repeat-containing double-stranded plasmid template with a primer that anneals up-or downstream of the repeats. The polymerization reaction was then carried out at 72°C in the presence of 3 mM Mg^2+^ as described in Materials and Methods. Note that upon denaturing and quick primer annealing, the plasmid template becomes a coil of intertwined single-stranded DNA segments, rather than a plain circular double-stranded DNA (Figure 1B). When the (A_2_G_3_)_10_ run serves as the template, polymerization stalls profoundly at its center (at the fifth and sixth repeats) and is unable to progress further in almost all templates (Figure 1A). In contrast, when the (C_3_T_2_)_10_ serves as the template, polymerization progresses through the repeats smoothly (Figure 1A). For the nonpathogenic repeats, when the (A_4_G)_10_ run is in the template, polymerization only mildly stalls at the sixth and seventh repeats, while the majority of DNA polymerases progress through the repeats. The pyrimidine (CT_4_)_n_ run in the template does not pose an obstacle for the DNA polymerase (Figure 1A).

**Figure 1.**
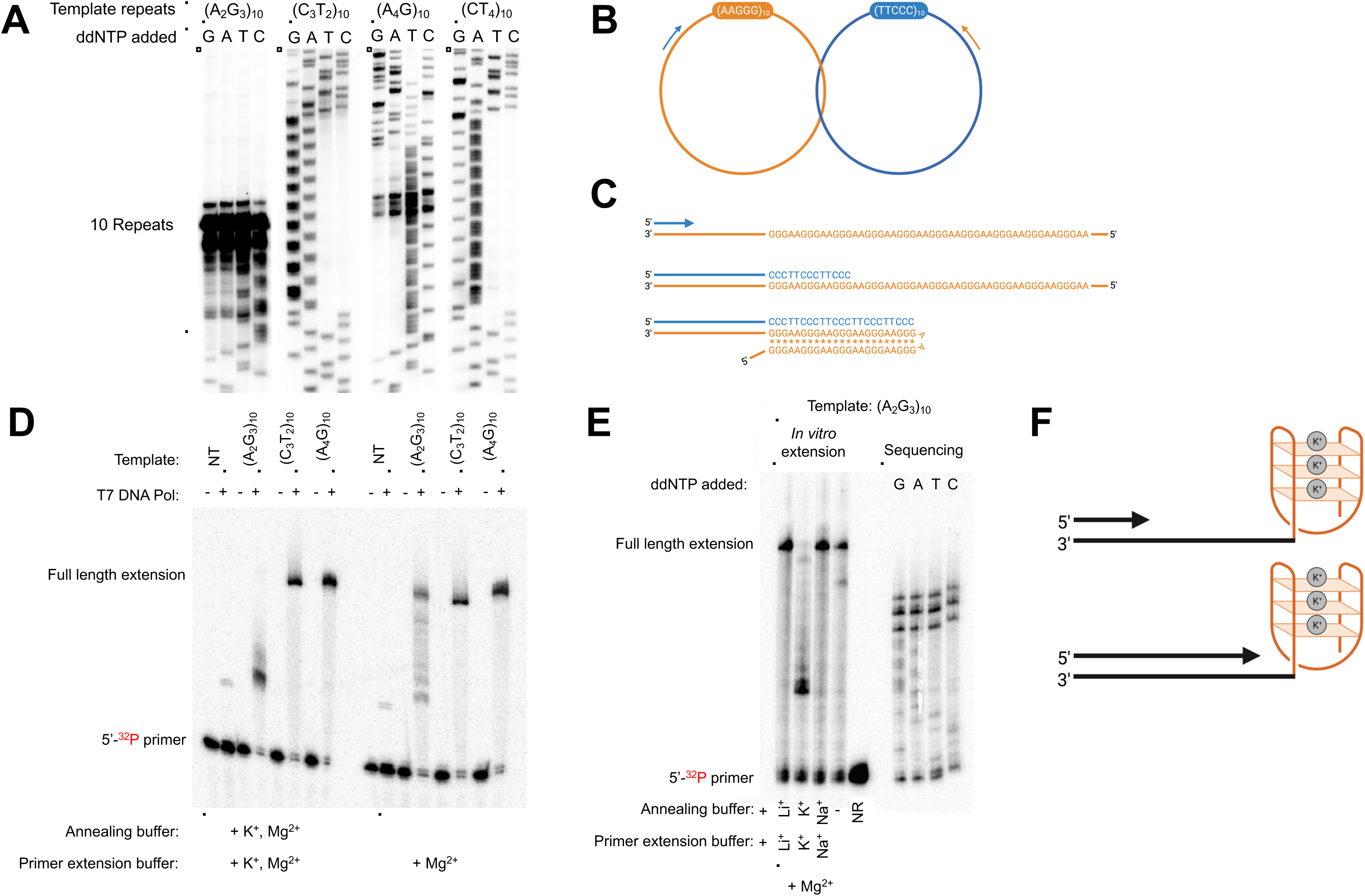
*In vitro* polymerization through pathogenic and nonpathogenic repeats. (A) Polyacrylamide gel electrophoresis separation of ThermoSequenase sequencing reactions performed as described in Materials and Methods. Briefly, 5μg of each plasmid and 0.5pmol of primer pre-annealed and the USBio Thermo Sequenase Cycle Sequencing Kit’s 3’-dNTP internal label cycling sequencing instructions were followed with several modifications detailed in Materials and Methods. (B) Schematic of denatured double-stranded plasmid with primers annealed to allow for ThermoSequenase polymerization through the purine-or pyrimidine-rich strand of (A_2_G_3_)_10_ or (A_4_G)_10_ in the template strand. Primers are 98 base pairs or 75 base pairs away from repeats for the purine-rich or pyrimidine-rich template, respectively. Created with BioRender. (C) Model for triplex formation as polymerase progresses through the repeats with the purine-rich strand as the template. Created with BioRender. (D) Polyacrylamide gel electrophoresis separation of *in vitro* T7 DNA polymerase primer extension reaction. 3.125µM single-stranded oligonucleotides bearing either (A_2_G_3_)_10_ (JH267), (T_2_C_3_)_10_ (JH268), or (A_4_G)_10_ (JH269) repeats were pre-annealed with a 2.5 µM 5’-^32^P labeled primer (JH270) in 10 mM Tris-HCl, pH 7.5, 0.1 mM EDTA by incubation at 95°C for 5 minutes, followed by gradual return to room temperature. 50mM K^+^ and 10mM Mg^2+^ were added to the annealing buffer in the left panel. Primer extension reactions consisted of 10nM primer/template and 1µM T7 DNA polymerase in 40 mM Tris-HCl, pH 7.5, 5mM dithiothreitol (DTT), and 55mM K^+^ (left panel). Primer extension was initiated by addition of 10mM MgCl_2_, followed by incubation at room temperature for 1 minute, and quenching with formamide/EDTA loading dye. Primer extension products were resolved by electrophoresis on a denaturing 10% polyacrylamide gel. NT=no template (primer added without oligonucleotide template). (E) The same general protocol was followed as in (D), with the following changes: 50mM LiCl, KCl, or NaCl were added to the annealing buffer and 55 mM of the indicated monovalent metal chloride salt was added to the primer extension buffer. Sequencing alongside primer extension reactions were conducted with the templates and primers used for the primer extension reactions and followed the protocol detailed in the Materials and Methods for the sequencing reactions in Figure 1A. NR=no reaction (no addition of MgCl_2_). (F) Model for G-quadruplex formation as T7 DNA polymerase progresses along the purine-rich template in the presence of K^+^ in the annealing and primer extension buffer.

Strikingly, for the relatively short (A_2_G_3_)_10_ template, a potent ThermoSequenase is basically halted at 72°C. The only instance when DNA polymerization was able to progress through the repeat under reaction conditions similar to ^21^: using Vent DNA polymerase at the extension temperature of 85°C and decreased to 1mM Mg^2+^ concentration (Figure S1A). This is indicative of the formation of an extremely stable secondary structure during the polymerization process.

*A priori* such a structure could either be (1) an H-r DNA triplex formed when DNA polymerase reaches the center of the template, or (2) a G-quadruplex formed by the G-rich template strand. The observed polymerase stalling at the center of the (A_2_G_3_)_10_ strand combined with the Mg^2+^-dependence of the stalling strongly implicates H-r DNA triplex formation, which, given the repeat’s base composition, would be among the strongest triplexes possible (Figure 1C). The lack of a potassium cation due to optimal ThermoSequenase reaction conditions and the presence of magnesium in our reaction renders G-quadruplex formation less likely ^37, 38^. Furthermore, G-quadruplex-forming sequences stall the polymerase directly at the 3’ end of the sequence ^38, 39^, rather than at the center. The minor stall within the nonpathogenic (A_4_G)_10_ repeat template is likely caused by a much weaker H-r DNA triplex. It was indeed found that A-rich H-r triplexes are stabilized by Zn^2+^, rather than Mg^2+^ cations ^40^.

We then investigated DNA polymerization through single-stranded CANVAS repeat-containing templates by bacteriophage T7 DNA polymerase. T7 DNA polymerase is a robust mesophilic enzyme that is often used as an *in vitro* model system for processive DNA polymerases and replication forks ^41^. Differently from ThermoSequenase, it is active at 25°C in the presence of both K^+^ and Mg^2+^ ions. For this initial experiment, we annealed a primer to a single-stranded template upstream of 10 repeat units: either (A_2_G_3_)_10_, (C_3_T_2_)_10_, or (A_4_G)_10_, followed by DNA polymerization. T7 DNA polymerase only stalls when (A_2_G_3_)_10_ serves as the template strand and does not stall when (C_3_T_2_)_10_ or the nonpathogenic repeat serve as the template strand (Figure 1D). Notably when both potassium and magnesium are present in the annealing and primer extension buffers, the stall is at the beginning of the repeats, while in the lack of potassium, DNA polymerase progresses through, but multiple weaker stalls are observed inside the repeat. Altogether, this is indicative of G-quadruplex formation stabilized by potassium ions.

Altering the annealing and primer extension buffer conditions to include various monovalent ions revealed that the polymerase stalling pattern with (A_2_G_3_)_10_ in the template strand changes depending on the surrounding ion concentration (Figure 1E). A significant stall occurs at the first repeat in the presence of potassium, while a weaker stall occurs just after half of the repeats when no additional salt is added to the annealing buffer (Figure 1E). The stall seen in high [K^+^] at the beginning of the repeats is likely caused by a G-quadruplex (Figure 1E, F). Meanwhile, the stall just after halfway through the repeats that occurs with no additional ions is likely an H-r triplex formed in a similar manner as the ThermoSequenase stall we observed. No stalling is observed when (C_3_T_2_)_10_ or (A_4_G)_10_ is in the template strand, regardless of surrounding ions (Figure S1B).

Altogether we conclude that depending on exact ion concentration, either a G-quadruplex or H-r triplex formed during polymerization through the (A_2_G_3_)_10_ template blocks DNA polymerase progression *in vitro*.

### (A_2_G_3_)_n_ repeats form triplex DNA in supercoiled DNA

While both the pathogenic (A_2_G_3_)_n_ and nonpathogenic (A_4_G)_n_ repeats are hPu/hPy mirror repeats, the former is G-rich and the latter is A-rich. *A priori*, both can form triplex H-DNA, but the (A_2_G_3_)_n_ repeat would most likely form an H-r triplex structure at physiological pH given the cytosine protonation required for the H-y isoform, while the (A_4_G)_n_ repeat could form either the H-r or H-y isoform, if it is able to form a triplex ^19, 42^. In addition, the (A_2_G_3_)_n_ repeat has the potential to form G4-DNA as it has regularly spaced G3 blocks. The (A_4_G)_n_ repeat, in contrast, can convert into the so-called propeller DNA (P-DNA) ^43^ or be a DNA unwinding element (DUE) as it has multiple A_n_-runs ^43, 44^.

To distinguish which of these structures are formed by those repeats in supercoiled DNA, we mapped single-stranded portions within the repeats *in vitro*. We used potassium permanganate (KMnO_4_), which preferentially modifies single-stranded thymines ^45^, thereby blocking Watson-Crick hydrogen bonding with a complementary strand and allowing their detection as polymerization termination sites in a primer extension assay. This approach could not be used for the (A_2_G_3_)_n_ strand since it stalls the polymerase, but interrogating the (C_3_T_2_)_n_ strand would allow us to distinguish between the candidate structures.

Upon potassium permanganate modification of plasmids with (A_2_G_3_)_10_ repeats, ThermoSequenase terminates in the wide area between the sixth and ninth repeats (Figure 2A). Meanwhile, the unmodified plasmid shows an almost undetectable level of polymerase termination (Figure S2). This data shows that the 5’-half of the polypyrimidine strand of the pathogenic repeat is single-stranded (Figure 2A), consistent with the H-r3 DNA triplex ^19^ (Figure 2B). At the same time, KMnO_4_ modification of the (A_4_G)_10_ repeat in supercoiled DNA shows evident termination signals at the beginning of the repeat combined with weak termination signals throughout the repeat (Figure 2A). This pattern is a stark contrast from the pathogenic repeat and suggests the (A_4_G)_10_ repeats in supercoiled DNA are transiently unwound, which is consistent with DUE chemical modification ^44^ (Figure 2C).

**Figure 2.**
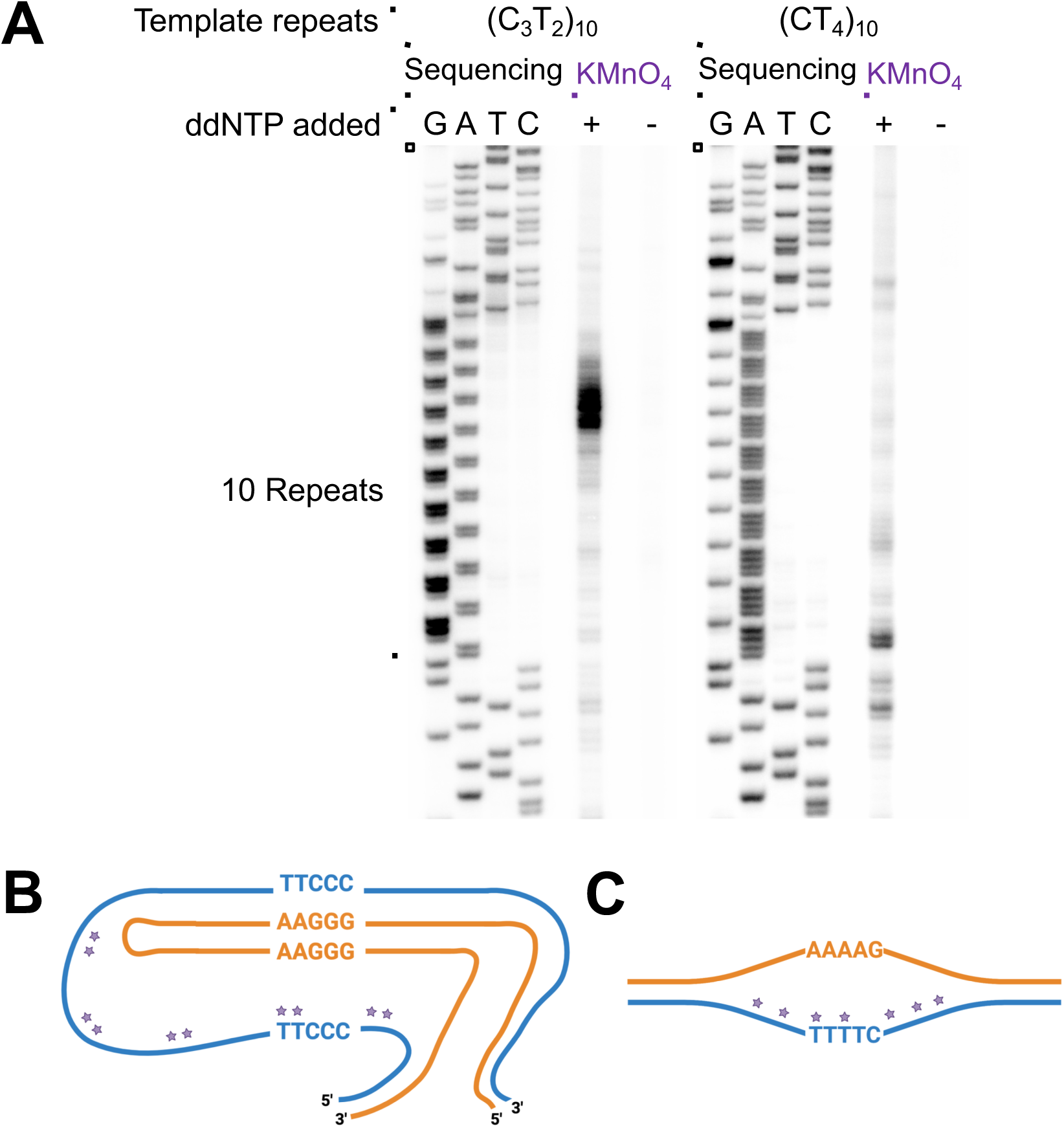
Potassium permanganate probing of pathogenic and nonpathogenic repeats. (A) Polyacrylamide gel electrophoresis separation of sequencing reactions and primer extension reactions on potassium permanganate or water treated repeat-containing plasmids using the pyrimidine-rich template. Repeat-containing supercoiled DNA was incubated with potassium permanganate or water, the DNA was precipitated, and used for a primer extension reaction as described in Materials and Methods. (B) H-r3 triplex predicted from chemical probing for (A_2_G_3_)_10_ repeats. (C) DNA unwinding element (DUE) predicted from chemical probing for (A_4_G)_10_ repeats. Purple stars in (B) and (C) represent possible KMnO_4_ modification sites.

### (A_2_G_3_)_n_ repeats have a higher propensity to form triplex DNA *in vivo* than (A_4_G)_n_ repeats

To examine the triplex-forming potential of (A_2_G_3_)_n_ and (A_4_G)_n_ repeats genome-wide *in vivo*, we reanalyzed an S1-END-seq dataset from a previous study^23^ which detected all triplex structures genome-wide in human cells. The S1-END-seq technique uses S1 nuclease to convert single-stranded DNA to double-strand breaks, which are then used as substrates for the attachment of high-throughput sequencing adapters ^23^. Triplex-forming regions were highlighted by this technique due to the presence of the extensive single-stranded regions in triplex H-DNA ^23^. Thus, to evaluate the triplex-forming potential of (A_2_G_3_)_n_ and (A_4_G)_n_ repeats, we compared the genomic coordinates of all such repeats in the human genome to the coordinates of peak regions identified by S1-END-seq. For both (A_2_G_3_)_n_ and (A_4_G)_n_ repeats, we see frequent overlap with the S1-END-seq peaks, growing to as much as 90% for very long repeat tracts, while overlap with randomly-generated genomic coordinates is in line with the 1.3% of the genome contained within peaks (Figure 3A). In contrast, G4-DNA motifs are not highly enriched in S1-END-seq peaks, with only 2-3% appearing in peaks regardless of motif length (Figure S3A). Other non-B-forming motifs are similarly not enriched in the S1-END-seq peaks (Figure S3B), with the sole exception of (AT)_n_ repeats (Figure S3C), demonstrating that the S1-END-seq assay is primarily specific for triplex-forming DNA. Thus, it is highly likely that both (A_2_G_3_)_n_ and (A_4_G)_n_ repeats form DNA triplexes *in vivo*. Note that these data cannot rule out G-quadruplex formation by the pathogenic repeats since S1-END-seq does not detect these structures as readily.

**Figure 3.**
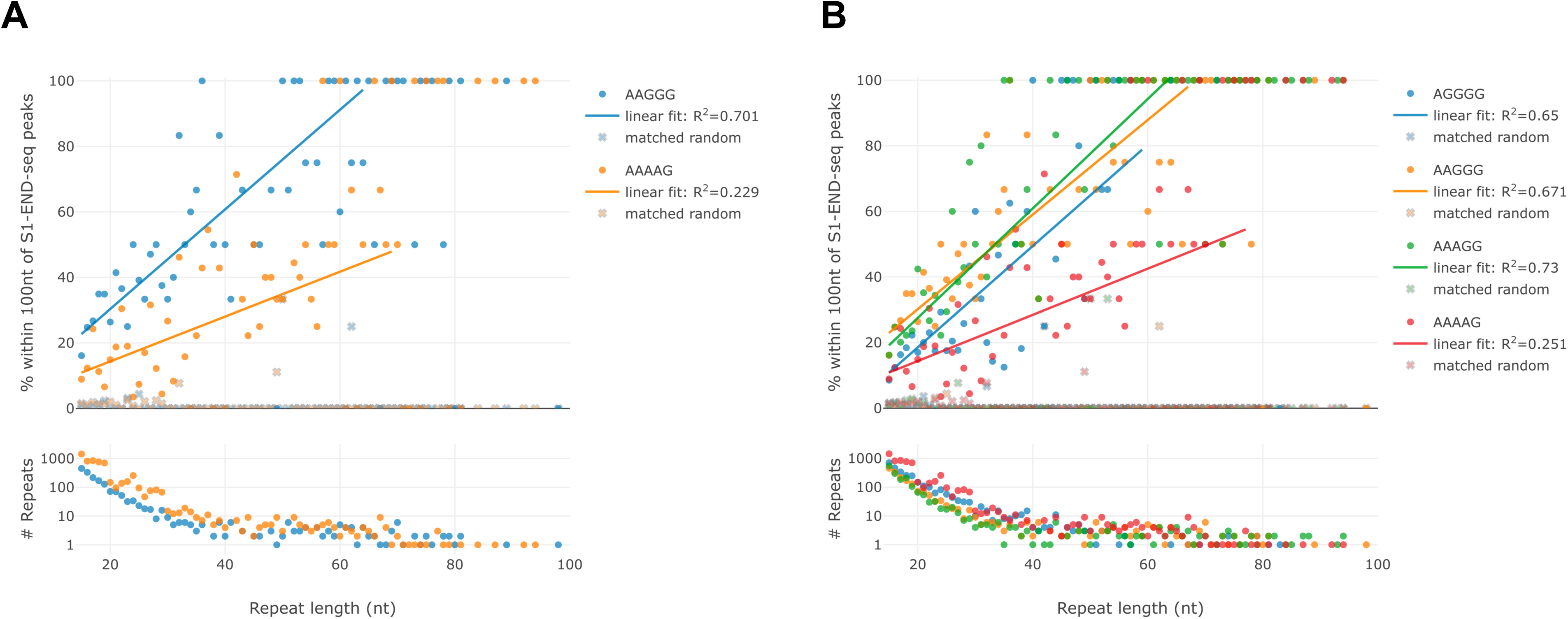
Bioinformatic analysis of S1-END-seq peaks to determine the triplex-forming potential of various repeats using data from ^23^. For each graph: Top: Graph depicting the percentage of repeats found within 100 nucleotides of S1-END-seq peaks as the repeat length increases. Bottom: Graph depicting the number of repeats as the repeat length increases. (A) Comparison of pathogenic (A_2_G_3_)_n_ and (A_4_G)_n_ motifs genome-wide. (B) Comparison of pentanucleotide motifs with increasing guanine:adenine ratio genome-wide.

Strikingly, (A_2_G_3_)_n_ repeats demonstrate consistently higher *in vivo* triplex-forming potential than (A_4_G)_n_ repeats along the axis of repeat length (Wilcoxon 1-sided test, p=2.7x10^-7^) (Figure 3A). Furthermore, other repeats known to be pathogenic – (AG_4_)_n_ and (A_3_G_2_)_n_ – also demonstrate higher *in vivo* triplex-forming potential than (A_4_G)_n_ repeats, all displaying a similar trend as (A_2_G_3_)_n_ (Figure 3B). Looking at a variety of hPu/hPy motifs with increasing numbers of adenines in a row, we see a pattern emerging, in which four or more adenines in a row becomes detrimental to triplex stability (Figure S3D). Looking at hexanucleotide motifs, though power-limited, we see that (A_4_G_2_)_n_ and (A_5_G_1_)_n_ show few signs of triplex formation, unlike all other hPu/hPy hexanucleotides (Figure S3E). Though (A_4_G_2_)_n_, (A_3_GAG)_n_ and (A_2_G)_n_ motifs all contain the same A:G ratio, only (A_4_G_2_)_n_ motifs are inhibited in triplex formation (Figure S3F). We hypothesize that the presence of longer adenine runs in double-stranded DNA favors stiff propeller DNA (P-DNA) ^43^, making triplex nucleation problematic. (A)_n_ repeats clearly do not form triplexes *in vivo*, while (G)_n_ repeats show a slight elevation in overlaps with S1-END-seq peaks (Figure S3G), consistent with other G4-DNA motifs.

### Pathogenic (A_2_G_3_)_n_ repeats stall replication in yeast in an orientation-dependent manner

Our *in vitro* polymerization through the repeats suggests stable H-r triplex or G-quadruplex formation as the polymerase progresses through the pathogenic repeats. It is not clear which structure would prevail during DNA replication *in vivo*, given two major factors: (1) far more components are at play in the replication fork as compared to DNA polymerase alone and (2) intranuclear conditions, including chromatin, DNA supercoiling, and ion concentrations, differ from that in the polymerization reactions. We hypothesized, based on our chemical probing and bioinformatic analysis, that the pathogenic (A_2_G_3_)_n_ repeats form H-r triplex DNA *in vivo* and would therefore stall replication only when the purine-rich strand resides on the lagging strand template. We also hypothesized that the nonpathogenic (A_4_G)_n_ repeats would not stall the replication fork to the same extent as the pathogenic repeats.

To study whether the expandable CANVAS repeats stall DNA replication *in vivo*, we cloned a longer (A_2_G_3_)_60_ repeat tract into the multicopy yeast pRS425 plasmid in both orientations with respect to the replication direction and analyzed their replication using 2-D electrophoretic analysis of replication intermediates (Figure 4) as described in ^46^.

**Figure 4.**
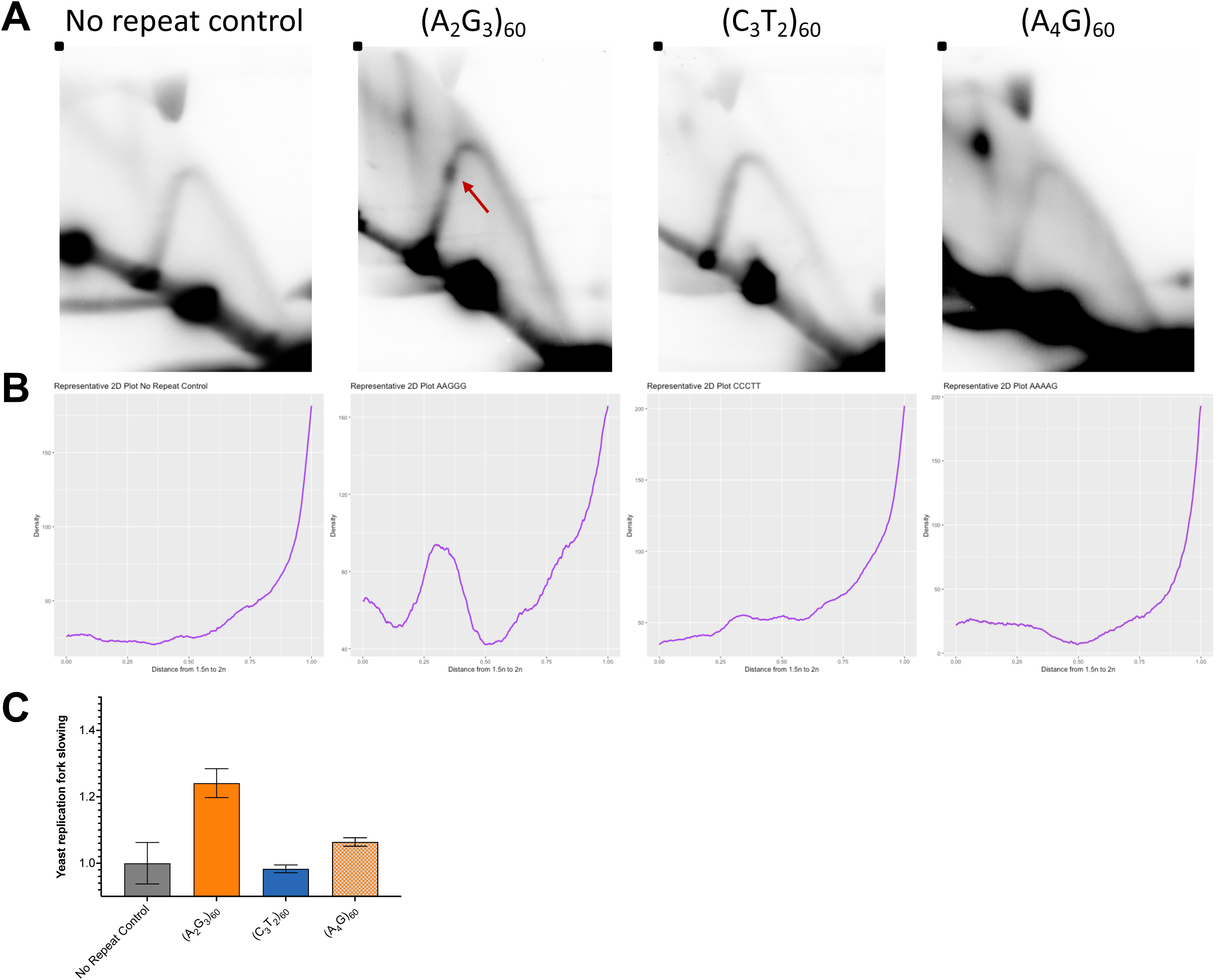
Analysis of yeast replication intermediates using two-dimensional gel electrophoresis. (A) Representative gels of the no repeat control, (A_2_G_3_)_60_, (C_3_T_2_)_60_, and (A_4_G)_60_ in the lagging strand template of replication in yeast from the yeast 2µ origin of replication. The red arrow indicates replication fork stalling. (B) Densitometry profiles along the arc starting at the 1.5n spot to the 2n spot. These profiles were used for quantification, which was determined as described in ^25, 46^. (C) Quantification of replication fork slowing via area analysis with the fold change increased normalized to the no repeat control arc. Error bars represent standard error of the mean and non-overlapping error bars were used to determine significance. Created with BioRender and prism.

There is no observable fork stall in the no repeat control plasmid (Figure 4 A,B). At the same time, the pathogenic repeat stalls replication fork progression only when its homopurine run is in the lagging strand template (Figure 4 A,B), as is evident from the presence of a bulge on the otherwise smooth descending half of the Y-arc. The presence of the (C_3_T_2_)_60_ run in the lagging strand template does not result in a defined stall site, but rather leads to a slight widening of the Y-arc downstream from the repeat. While the reasons for this arc widening are not clear, it might indicate minor slowing of the replication fork progression as it passes through the repeat (Figure 4 A,B). Regardless, quantification of the stalling (Figure 4 B,C) reveals a significant increase in replication fork stalling with (A_2_G_3_)_60_ in the lagging strand template over the no repeat control and the flipped orientation (Figure 4C). Similar orientation-dependence was previously observed for the Friedreich’s ataxia (GAA)_n_ repeat in many systems ^33, 47, 48^, which forms H-r triplexes *in vitro* ^49–52^ and *in vivo* ^23^, suggesting the pathogenic CANVAS repeats may also form an H-r triplex *in vivo*. Importantly, the replication fork does not stall significantly more than the no repeat control when (A_4_G)_60_ is in the lagging strand template, in line with our hypothesis based on little *in vitro* polymerase stalling at the (A_4_G)_10_ repeats (Figure 1) and the lack of structure formation seen on (A_4_G)_10_ chemical probing (Figure 2). Altogether, these *in vivo* data mirror our *in vitro* results in their orientation-dependence and pathogenic repeat-dependence.

### Pathogenic (A_2_G_3_)_n_ repeats stall replication in human cells in an orientation-dependent manner

Does the replication fork stalling by (A_2_G_3_)_n_ repeats in yeast hold true in human cells? To answer this question, we utilized an episome replicating in human HEK293T cells described by us earlier ^26^. Briefly, this episome contains both the SV40 origin of replication and the T-antigen driving its replication initiation and elongation. Thus, the more it replicates, the more T-antigen is produced, driving subsequent rounds of replication. As a result, this system generates a high amount of replication intermediates, making their electrophoretic analysis feasible and relatively easy. The (A_2_G_3_)_60_ repeat was cloned into this plasmid in two orientations relative to the SV40 origin.

Replication fork stalling patterns caused by the pathogenic (A_2_G_3_)_n_ repeats in human cells appear to be fundamentally similar to that in yeast (Figure 5). When in the lagging strand template, the (A_2_G_3_)_60_ run causes a very prominent replication stall on the ascending half of the Y-arc, while no stall is detected in the no repeat control (Figure 5A,B). In the opposite orientation, the (C_3_T_2_)_60_ run on the lagging strand template does not cause prominent fork stalling (Figure 5A,B). Also similar to the yeast 2D data, the non-pathogenic (A_4_G)_n_ repeat does not cause significant fork stalling (Figure 5).

**Figure 5.**
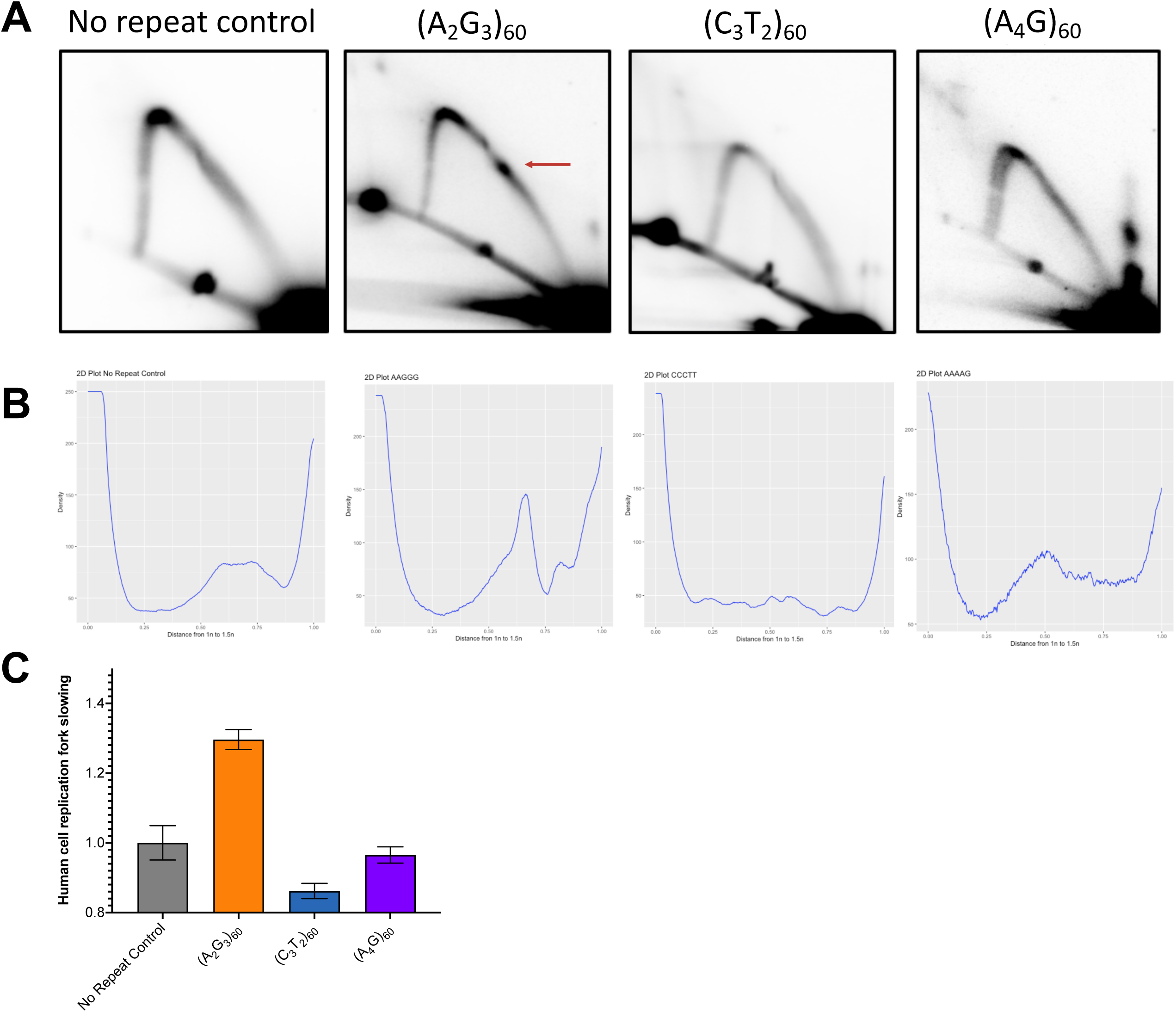
Analysis of human cell replication intermediates using two-dimensional gel electrophoresis. (A) Representative gels of the no repeat control, (A_2_G_3_)_60_, (C_3_T_2_)_60_, and (A_4_G)_60_ in the lagging strand template of replication from the SV40 origin of replication. The red arrow indicates replication fork stalling. (B) Densitometry profiles along the arc starting at the 1n spot to the 1.5n spot. These profiles were used for quantification, which was determined as described in ^25, 46^. (C) Quantification of replication fork slowing via area analysis with the fold change increased normalized to the no repeat control arc. Error bars represent standard error of the mean and non-overlapping error bars were used to determine significance. Created with BioRender and prism.

The consistent throughline in the *in vitro* and *in vivo* replication data is that the pathogenic (A_2_G_3_)_n_ repeats are an obstacle to DNA polymerases and replication forks particularly when the purine-rich strand is more single-stranded.

## DISCUSSION

CANVAS is a recently discovered neurodegenerative RED characterized by a spectrum of clinical manifestations, including but not limited to cerebellar ataxia, neuropathy, and vestibular areflexia ^1, 2^. Though it is estimated to be the most common cause of hereditary late-onset ataxia ^1, 4^, very little is known about its genetics and pathogenesis. An unusual feature of this RED is that a change in the sequence from the nonpathogenic (A_4_G)_11-100_ to the pathogenic (A_2_G_3_)_250-2,000_ causes CANVAS, rather than only the expansion of a repeat ^1^. The recent discovery of various pathogenic and benign repeat iterations poses several interesting questions: (1) Do the repeats form a non-B DNA structure that is integral to its instability, as other RED repeats do? (2) Is this non-B DNA structure the same for different pathogenic repeats? (3) Is structure formation by only the pathogenic repeats the underlying reason for their pathogenicity? (4) How do the repeats expand and cause disease?

No model systems have so far been developed to study CANVAS’s genetics or pathogenesis. Clues to RED pathogenesis often lie within the repeats themselves, the non-B structures they form, and their interactions with cellular machinery. For example, the (GAA)_exp_ repeats implicated in FRDA are at the heart of the expansion mechanism and disease pathogenesis: (GAA)_n_-related issues with replication and DNA repair contribute to the expansion of (GAA)_n_ repeats (reviewed in ^34^) and expanded (GAA)_n_ repeats block transcription, contributing to the loss of function pathogenesis of FRDA (reviewed in ^12^). Triplex formation of the (GAA)_exp_ allele has thereby been shown to be central to both FRDA genetics and pathogenesis. Therefore, it is crucial to study the pathogenic (A_2_G_3_)_n_ repeat in terms of its propensity to form a non-B DNA structure, ability to impede cellular machinery such as replication, and mechanisms of instability.

Using an arsenal of tools developed by our lab and others, we examined the CANVAS-causing (A_2_G_3_)_n_ repeats and established key DNA-centric characteristics that may help unravel how these repeats expand and cause disease.

Using probing with a chemical specific to single-stranded thymines, we found that (A_2_G_3_)_10_, but not (A_4_G)_10_, forms an H-r DNA triplex in supercoiled DNA (Figure 2A). Recent S1-END-seq and permanganate footprinting data has shown that many hPu/hPy mirror repeats form triplexes *in vivo* ^23, 53^. In fact, S1-END-seq indicated triplex formation in lymphoblasts derived from a (GAA)_exp_-harboring FRDA patient and not in an unaffected sibling ^23^. We suspect the same may be true for CANVAS patients and we wondered if we could glean information about the CANVAS repeats’ secondary structure by determining if these repeats form secondary structures elsewhere in the genome. Using the S1-END-seq data ^23^, we indeed found that long tracts of (A_2_G_3_)_n_ and (A_4_G)_n_ motifs overlap with S1-END-seq peaks genome-wide, indicative of their triplex formation *in vivo* (Figure 3). Remarkably, the propensity for triplex formation is higher for (A_2_G_3_)_n_ than (A_4_G)_n_ motifs *in vivo*, supporting our hypothesis that the pathogenic repeats form a more stable triplex than the nonpathogenic repeats, potentially leading to downstream repeat instability.

Transient non-B DNA structure formation by the repeats blocks DNA polymerization through the repeat at the center of the repeat *in vitro* in an orientation-dependent manner, *i.e.* when the homopurine run is on the template strand (Figure 1A). This is not the case for the nonpathogenic (A_4_G)_n_ repeat. Many triplex-forming repeats have been shown to stall DNA polymerases *in vitro* particularly strongly when the homopurine strand is the template strand ^21, 32, 54–57^. In fact, the pattern we observed *in vitro* with prominent stalling once the polymerase reaches halfway through the repeats is almost identical to that seen with another H-r triplex-forming sequence ^54, 55^. Interestingly, the pathogenic CANVAS repeat, when present in ssDNA, can adopt alternative structures depending on solution conditions. *In vitro* extension experiments using T7 DNA polymerase demonstrated stalling patterns that indicate the formation of G-quadruplex (Figure 1D,E). Solution conditions that stabilize G-quadruplexes result in a strong stall immediately upstream of the (A_2_G_3_)_10_ repeat. Ambient cellular conditions as well as additional replication proteins likely play a major role in determining the secondary structure formed by the CANVAS-causing repeats during replication *in vivo*.

Would such a potent repeat-mediated DNA polymerization block *in vitro* manifest itself *in vivo*? To study the effect of this repeat on replication in yeast, we used a repeat-bearing plasmid with a yeast 2µ origin of replication, which uses the host replisome to replicate through the repeats. For replication in human cells, we used a repeat-bearing plasmid with an SV40 origin of replication that expresses T antigen. Remarkably, the pathogenic (A_2_G_3_)_n_ repeat blocks DNA replication in both yeast and human cells in an orientation-dependent manner: when the (A_2_G_3_)_n_ run is in the lagging strand template (Figures 4 and 5).

H-motifs have been shown to stall replication when the homopurine strand is in the lagging strand template in bacteria ^58^, yeast ^33, 47, 48^, human cells, and human cell extracts ^32^. This replication fork stalling pattern is in stark contrast to orientation-independent hairpin-forming sequences ^59, 60^ while the pattern of replication fork stalling for G-quadruplexes is more complicated with evidence of G-quadruplexes impeding the replication fork when in the leading ^61–63^ or lagging ^64^ strand template. Often, a G-quadruplex stabilizer or the knockout of a G-quadruplex unwinder is necessary to cause stalling ^61, 64^. Therefore, the CANVAS repeat replication stalling pattern that is conserved from yeast to human cells is more consistent with a triplex-caused polymerization arrest *in vivo*, though G-quadruplex formation cannot be ruled out.

The dynamic nature of non-B DNA structure formation may allow for the formation of both a triplex and a G-quadruplex by the pathogenic repeats, depending cellular conditions. Experiments to determine genetic controls of this replication fork stalling are underway and will surely shed more light on the non-B DNA structure(s) involved in the orientation-dependent replication fork stalling observed at the pathogenic repeats.

Overall, we have found that the pathogenic (A_2_G_3_)_n_ repeat forms a triplex *in vitro* and stalls replication in an orientation-dependent manner *in vitro* and *in vivo*. In each experiment, we have juxtaposed the pathogenic (A_2_G_3_)_n_ repeat with the nonpathogenic (A_4_G)_n_ repeat, finding the nonpathogenic repeat largely behaves similar to a nonrepetitive sequence and thus does not impede replication to the extent that (A_2_G_3_)_n_ does. Therefore, we believe we have uncovered features of the pathogenic allele that are integral to its instability and pathogenicity. This illuminates an important next step: studying the genetics of CANVAS and how the repeats expand and cause disease. To this point, work to establish model systems to study the repeats’ instability (contraction and expansion) is currently underway and is a crucial start to understanding this recently characterized disease. Similarly, conducting these structure-functional analysis experiments with additional pathogenic and nonpathogenic alleles may illuminate a pattern suggestive of one structure over another, though it is possible different structures are formed by different repeats, leading to the same disease.

## Supporting information

Supplemental Materials for Hisey et al, 2023

## DATA AVAILABILITY

The data underlying this article are available in the article and in its online supplementary material.

## SUPPLEMENTARY DATA

Supplementary data is available in a separate PDF.

## ACKNOWLEDGEMENTS

We thank Catherine Freudenreich, Mitch McVey, Claire Moore, Ralph Scully and members of the Mirkin laboratory for their integral input to this project.

