## Supplemental Materials for Hisey et al, 2023 for "Pathogenic CANVAS (AAGGG)_n_ repeats stall DNA replication due to the formation of alternative DNA structures"

#### Supplemental Material

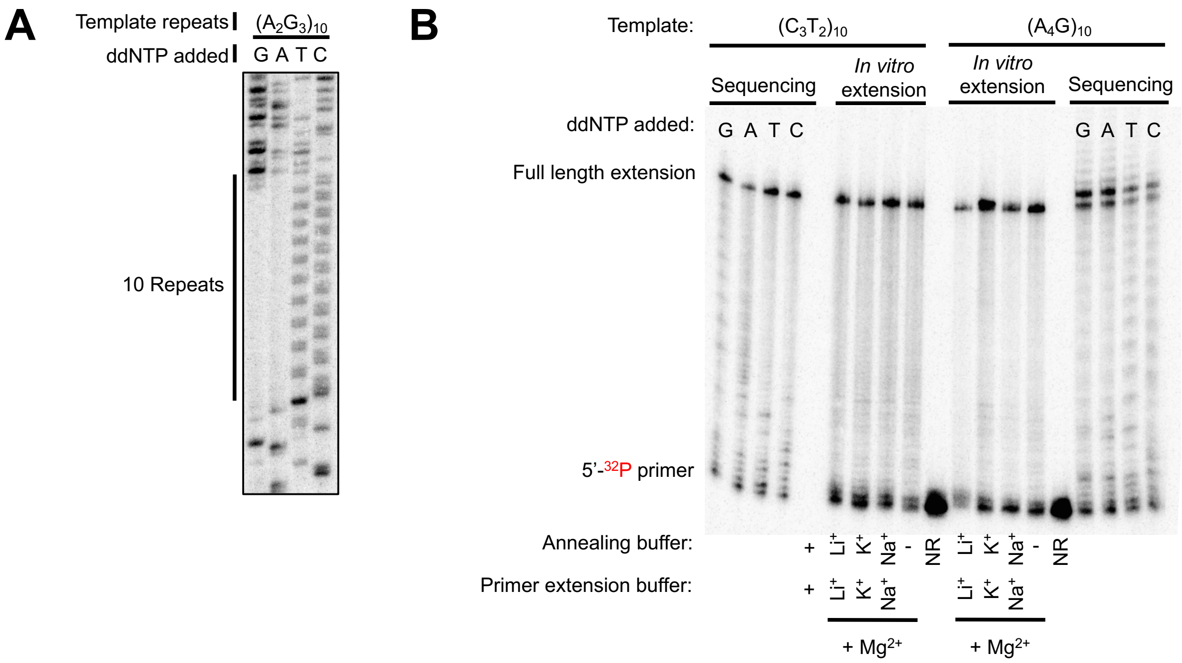

**Figure S1**. ***In vitro* polymerization with Vent polymerase and T7 DNA polymerase.** (A) *In vitro* polymerization through pathogenic repeats with Vent (exo-) polymerase. Polyacrylamide gel electrophoresis separation of sequencing reactions with Vent (exo-) polymerase with an 80°C extension temperature, using a plasmid template containing $(A_2G_3)_{10}$ repeats and a primer 98 base pairs away from repeats. (B) T7 DNA polymerase primer extension reactions as seen in Figure 1E with $(C_3T_2)_{10}$ and $(A_4G)_{10}$ in the template strand. Sequencing reactions serving as a ladder are to the right and left of T7 primer extension reactions for $(C_3T_2)_{10}$ and $(A_4G)_{10}$ primer extension reactions, respectively.

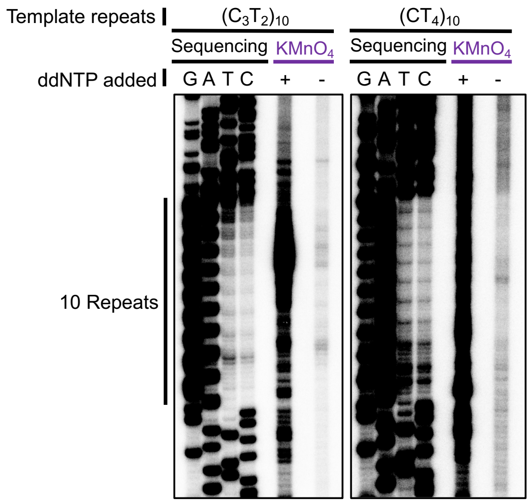

**Figure S2.** The same polyacrylamide gels seen in Figure 2A with increased contrast to visualize the *in vitro* chemical probing control with water instead of potassium permanganate.

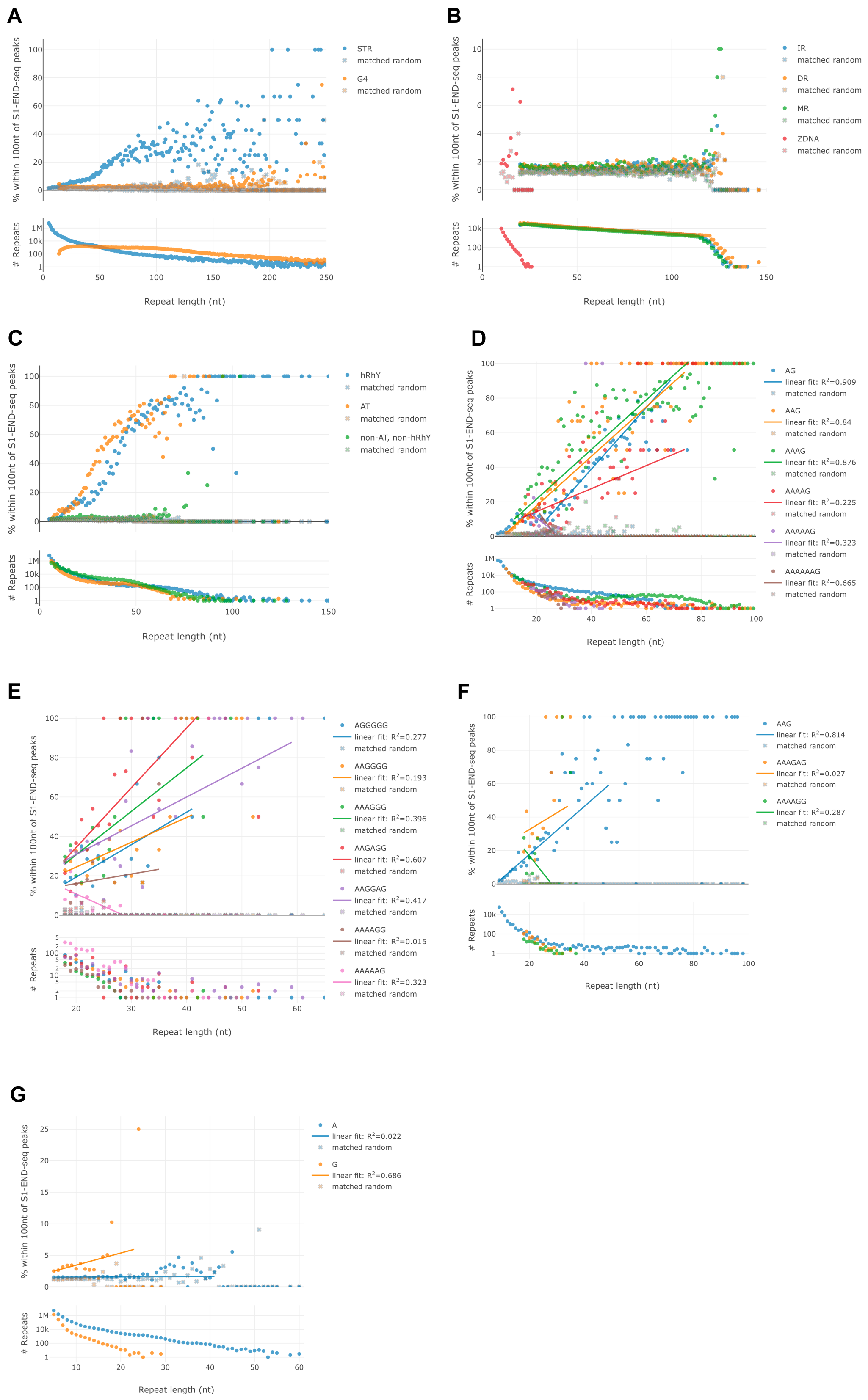
A
% within 100nt of S1-END-seq peaks
STR
matched random
G4
matched random
### Repeats
Repeat length (nt)
B
IR
matched random
DR
matched random
MR
matched random
ZDNA
matched random
C
hRhY
matched random
AT
matched random
non-AT, non-hRhY
matched random
D
AG
linear fit: $R^2$=0.909
matched random
AAG
linear fit: $R^2$=0.84
matched random
AAAG
linear fit: $R^2$=0.876
matched random
AAAAG
linear fit: $R^2$=0.225
matched random
AAAAAG
linear fit: $R^2$=0.323
matched random
AAAAAAG
linear fit: $R^2$=0.665
matched random
E
AGGGGG
linear fit: $R^2$=0.277
matched random
AAGGGG
linear fit: $R^2$=0.193
matched random
AAAGGG
linear fit: $R^2$=0.396
matched random
AAGAGG
linear fit: $R^2$=0.607
matched random
AAGGAG
linear fit: $R^2$=0.417
matched random
AAAAGG
linear fit: $R^2$=0.015
matched random
AAAAAG
linear fit: $R^2$=0.323
matched random
F
AAG
linear fit: $R^2$=0.814
matched random
AAAGAG
linear fit: $R^2$=0.027
matched random
AAAAGG
linear fit: $R^2$=0.287
matched random
G
A
linear fit: $R^2$=0.022
matched random
G
linear fit: $R^2$=0.686
matched random

**Figure S3.** Bioinformatic analysis of S1-END-seq peaks to determine the triplex-forming potential of various repeats using data from [1]. For each graph: Top: Graph depicting the percentage of repeats found within 100 nucleotides of S1-END-seq peaks as the repeat length increases. Bottom: Graph depicting the number of repeats as the repeat length increases. (A) Comparison of STR and G4 motifs. (B) Comparison of various non-B motifs. (C) Comparison of homopurine/homopyrimidine motifs, $(AT)_n$ repeats, and non-AT, non-homopurine/homopyrimidine motifs. (D) Comparison of homopurine/homopyrimidine motifs with increasing numbers of adenines in a row. (E) Comparison of homopurine/homopyrimidine hexanucleotide motifs. (F) Comparison of homopurine/homopyrimidine motifs with the same ratio of adenines to guanines. (G) Comparison of $(A)_n$ and $(G)_n$ motifs.

**Construction of plasmids**

Yeast 2D*:* Primers JH92 and JH93 were used to amplify $(A_2G_3)_{60}$ from pJH7 to add BsrGI restriction sites to the ends of this PCR product and this product was inserted into plasmid pRS425-UIRLB [2] by digestion with BsrGI and ligation. Clones identified as correct using (1) PCRs flanking the ligation integration site and (2) Repeat PCR protocol Taq 1 were sent for sequencing and the correct clones were named pJH1 (purine lagging strand template) and pJH26 (pyrimidine lagging strand template). Plasmid pJH4 with the $(A_2G_3)_{60}$ repeats replaced by $(A_4G)_{60}$ was ordered from Genescript.

Human cell 2D: To construct a plasmid to observe replication through $(A_2G_3)_n$ repeats in the lagging strand template orientation in human cells using two-dimensional gel electrophoresis, we replaced an existing plasmid with $(GAA)_{100}$ repeats (pJH29) with $(A_2G_3)_{60}$ repeats from pJH19 (ordered from Genescript) using restriction sites PspXI and Esp3I. Correctly ligated clones were screened using restriction digests of restriction enzyme sites within the vector and insert. Cloning junctions of correct clones were sequenced, repeat lengths were checked using PCR program Taq 1, and the correct plasmid was named pJH5. To flip the cassette, we amplified UR-$(A_2G_3)_{60}$-A3 cassette with primers JH295 and JH296 that incorporate restriction enzyme sites BmgI and BglII on either side of the repeats; these restriction sites were used to insert $(C_3T_2)_{60}$ into pJH5. Successfully ligated colonies were screened with two PCRs flanking the integration sites: (1) NHM30 and JK149 and (2) NHM29 and NHM27. Cloning junctions of correct clones were sequenced, repeat lengths were checked using PCR program Taq 1, and the correct plasmid was named pJH5. Gibson assembly with (1) Primers JH53 and JH54 and plasmid pJH4 and (2) Primers JH55 and JH56 and plasmid pJH5 was used to replace $(A_2G_3)_{60}$ in pJH5 with $(A_4G)_{60}$ from plasmid pJH4. Cloning junctions of correct clones were sequenced, repeat lengths were checked using PCR program Taq 2, and the correct plasmid was named pJH11.

**Analysis of two-dimensional gel electrophoresis.**

Two-dimensional gel electrophoresis quantification was carried out as described in [2] and demonstrated visually in Figure 4 of [3]. For each biological replicate, of which there were three for yeast and three for human cell replication, the following was performed. Using ImageJ, the Y-arc was traced from the 1.5n spot to the 1n spot for yeast 2Ds and from the 1n spot to the 1.5n spot for human cell 2Ds in order to have a measure of the density of the arc. The same was done for a line above and below the arc to have an average background signal. A line was drawn to estimate where a smooth arc profile would be, as in [3]. Using ImageJ, the area of the stall (1 in Figure 4 of [3]) was added to the area representing the background (2 in Figure 4 of [3]) and divided by the area of the background (2 in Figure 4 of [3]). This was normalized to that of the no repeat control for both yeast and human cell 2D gels. These values are on the quantification bar graphs in Figures 4 and 5.

**Supplemental Table 1. Standard Phusion PCR Program**

| **Step** | **Temperature** | **Time** |
|---|---|---|
| Initial denaturation | 98C | 30s |
| Cycle denaturation | 98C | 15s |
| Cycle annealing | Calculated using NEB Tm calculator | 50s |
| Cycle extension | 72C | 30s/kb |
| X35 cycles | | |
| Final extension | 72C | 5m |

| | | |
|---|---|---|
| | 4C | 5m |
| | 12C | Infinity |

**Supplemental Table 2. Specific Phusion PCR programs**

| Purpose | Primers | Primer Annealing | Product size (bp) | Extension Time and Temp if different | Template |
|---|---|---|---|---|---|
| $(A_2G_3)_{60}$ yeast 2D fragment | JH92, JH93 | 72C | 460 | 80C, 5 minutes | pJH7 |
| Amplify $(A_4G)_{60}$ in yeast 2D | JH9, JH10 | 72C | 439 | 75C, 5 min, x5 cycles 75C, 3 min, x30 cycles | pJH11 and pJH9 |

**Supplemental Table 3. Specific Taq PCR programs for checking $A_2G_3$ and $A_4G$ repeat length**

| Program name | Purpose | Primers | Denature | Annealing | Product size | Extension Time and Temp if different |
|---|---|---|---|---|---|---|
| Taq 1 | Amplify $(A_2G_3)_{60}$ within URA3 intron | JH9 JH10 | Initial: 95C, 2m Cycles: 95C, 1m | 78C, 2m | 439bp | 5 cycles: 80C, 5m 30 cycles: 80C, 3m |
| Taq 2 | Amplify $(A_4G)_{60}$ in human cell 2D plasmid | RM298 RM299 | Initial: 95C, 2m Cycles: 95C, 1m | 63C, 2m | 846bp | 5 cycles: 80C, 5m 30 cycles: 80C, 3m |

**Supplemental Table 4. Plasmids used in this study**

| Plasmid name | E. coli Stock | Notes |
|---|---|---|
| pJH2 | JAH104 | Plasmid for in vitro polymerization with $(A_2G_3)_{10}$ repeats |
| pJH9 | JAH112 | Plasmid for in vitro polymerization with $(A_4G)_{10}$ repeats |
| pJH7_UR-$(A_2G_3)_{60}$-A3 TRP1_orientation1 | JAH7 | Plasmid to amplify $(A_2G_3)_{60}$ for yeast 2D plasmid construction |

| | | |
|---|---|---|
| pJH1 | JAH69 | Plasmid for yeast 2D with $(A_2G_3)_{60}$ repeats in lagging strand template |
| pJH26 | JAH65 | Plasmid for yeast 2D with $(C_3T_2)_{60}$ repeats in lagging strand template |
| pJH4 | JAH108 | Plasmid for yeast 2D with $(A_4G)_{60}$ repeats in lagging strand template |
| pJH27 | JAH83 | Plasmid for yeast 2D with a no repeat control sequence |
| pJH29 | | Plasmid able to replicate in human and yeast with $(GAA)_{100}$ repeats |
| pJH19 | JAH1 | Plasmid ordered from genescript with $(A_2G_3)_{60}$ and pieces of surrounding URA3 intron and gene |
| pJH5_$A_2G_3$_60_mammalianrep | JAH29 | Plasmid able to replicate in human and yeast with $(A_2G_3)_{60}$ in the lagging strand template of replication |
| pJH6_$C_3T_2$_60_mammalianrep | MC5 | Plasmid able to replicate in human and yeast with $(C_3T_2)_{60}$ in the lagging strand template of replication |
| pJH11_$A_4G$_mammalianrep | JAH130 | Plasmid able to replicate in human and yeast with $(A_4G)_{60}$ in the lagging strand template of replication |
| pJH28 | | No repeat control plasmid for UR-$(A_2G_3)_{60}$-A3 cassette for arm loss assay |
| pJH17 UAS-$(A_2G_3)_{84}$-Pgal-CAN1 | JAH41 | Plasmid containing $(A_2G_3)_{84}$ replacing $(CAG)_{140}$ from Kim *et al.* 2017 [4] |

**Supplemental Table 5. Primers used in this study**

| **Primer Name** | **Primer Sequence** | **Notes** |
|---|---|---|
| Repeat amplification primers | | |
| JH9_A2G3repeatslong F2 | GCACGGTCCCAATTCTGCAGATATCCATCACACTGGCGGCCGCTCGAGTGCAGACCTCAAATTCG | To amplify for 439bp product to check $(A_2G_3)_{60}$ repeats for yeast 2D, |

| | | |
|---|---|---|
| JH10_A2G3repeatslong R2 | AGGTTGAGGCCGTTGAGCACCGCCGCCGCAAGGAATGGTGCATGCTCGATGTGCAGAACCTGAAGCTTG | mammalian 2D, and arm loss assay |
| Sequencing and chemical probing | | |
| JH271_C_template_dATP_lab | AGCGCTATATGCGTTGATGC | Pyrimidine repeat template strand |
| JH272_G_template_dATP_lab | GCTTAAAAAGATTCTCTTTTTTTATGATATTTGTACAT | Purine repeat template strand |
| Yeast 2D Plasmid Construction | | |
| JH92_A2G3_60_BsrGI_F | GCATACTGATGTACATCTGCAGATATCCATCACACTGGC | To amplify $(A_2G_3)_{60}$ from pJH7 to add BsrGI restriction sites and insert fragment into plasmid pRS425-UIRLB [2] |
| JH93_A2G3_60_BsrGI_R | GCATACTGATGTACATAGTAGGTTGAGGCCGTTGAGC | |
| T7 primer extension | | |
| JH267_bottom_A2G3 | GGGAAGGGAAGGGAAGGGAAGGGAAGGGAAGGGAAGGGAAGGGAAGGGAACAGTACTCCGCTCTATAGTTTGTATCGTCACCATAACTCTGTAACTAG | Templates for T7 primer extension |
| JH268_bottom_T2C3 | CCCTTCCCTTCCCTTCCCTTCCCTTCCCTTCCCTTCCCTTCCCTTCCCTTCAGTACTCCGCTCTATAGTTTGTATCGTCACCATAACTCTGTAACTAG | |
| JH269_bottom_A4G | GAAAAGAAAAGAAAAGAAAAGAAAAGAAAAGAAAAGAAAAGAAAAGAAAACAGTACTCCGCTCTATAGTTTGTATCGTCACCATAACTCTGTAACTAG | |
| JH270_Fork_Sub_F | CTAGTTACAGAGTTATGGTGACGATACAAACTAT | Primer for T7 extension |
| JH349_bottom_A2G3_10 | CTAGTTACAGAGTTATGGTGACGATACAAACTATAGAGCGGAGTACTGTTCCCTTCCCTTCCCTTCCCTTCCCTTCCCTTCCCTTCCCTTCCCTTCCCTTCCC | Ladder for A2G3 primer extension |
| JH350_bottom_A2G3_8 | CTAGTTACAGAGTTATGGTGACGATACAAACTATAGAGCGGAGTACTGTTCCCTTCCCT | |

| | | |
|---|---|---|
| | TCCCTTCCCTTCCCTTCCCTT CCCTTCCC | |
| JH351_bottom_A2G3_ 6 | CTAGTTACAGAGTTATGGTG ACGATACAAACTATAGAGC GGAGTACTGTTCCCTTCCCT TCCCTTCCCTTCCCTTCCC | |
| JH352_bottom_A2G3_ 4 | CTAGTTACAGAGTTATGGTG ACGATACAAACTATAGAGC GGAGTACTGTTCCCTTCCCT TCCCTTCCC | |
| JH353_bottom_A2G3_ 2 | CTAGTTACAGAGTTATGGTG ACGATACAAACTATAGAGC GGAGTACTGTTCCCTTCCC | |
| JH354_bottom_T2C3_ 10 | CTAGTTACAGAGTTATGGTG ACGATACAAACTATAGAGC GGAGTACTGAAGGGAAGGG AAGGGAAGGGAAGGGAAGG GAAGGGAAGGGAAGGGAAG GG | Ladder for T2C3 primer extension |
| JH355_bottom_T2C3_ 8 | CTAGTTACAGAGTTATGGTG ACGATACAAACTATAGAGC GGAGTACTGAAGGGAAGGG AAGGGAAGGGAAGGGAAGG GAAGGGAAGGG | |
| JH356_bottom_T2C3_ 6 | CTAGTTACAGAGTTATGGTG ACGATACAAACTATAGAGC GGAGTACTGAAGGGAAGGG AAGGGAAGGGAAGGGAAGG G | |
| JH357_bottom_T2C3_ 4 | CTAGTTACAGAGTTATGGTG ACGATACAAACTATAGAGC GGAGTACTGAAGGGAAGGG AAGGGAAGGG | |
| JH358_bottom_T2C3_ 2 | CTAGTTACAGAGTTATGGTG ACGATACAAACTATAGAGC GGAGTACTGAAGGGAAGGG | |
| JH359_bottom_A4G_1 0 | CTAGTTACAGAGTTATGGTG ACGATACAAACTATAGAGC GGAGTACTGTTTTCTTTTCTT TTCTTTTCTTTTCTTTTCTTTT CTTTTCTTTTCTTTTC | Ladder for A4G primer extension |
| JH360_bottom_ A4G _8 | CTAGTTACAGAGTTATGGTG ACGATACAAACTATAGAGC GGAGTACTGTTTTCTTTTCTT TTCTTTTCTTTTCTTTTCTTTT CTTTTC | |

| | | |
|---|---|---|
| JH361_bottom_ A4G _6 | CTAGTTACAGAGTTATGGTGACGATACAAACTATAGAGCGGAGTACTGTTTTCTTTTCTTTTCTTTTCTTTTCTTTTC | |
| JH362_bottom_ A4G _4 | CTAGTTACAGAGTTATGGTGACGATACAAACTATAGAGCGGAGTACTGTTTTCTTTTCTTTTCTTTTC | |
| JH363_bottom_ A4G _2 | CTAGTTACAGAGTTATGGTGACGATACAAACTATAGAGCGGAGTACTGTTTTCTTTTC | |
| Human Cell 2D Plasmid Construction and Checking | | |
| JH295_Fliprepeats-F-BglII-BmgBI | TTCTTAGACCCACGTCAGCTTTTCAATTCAATTCAT | To amplify for $(C_3T_2)_{60}$ human 2D plasmid |
| JH296_Fliprepeats-R-BglII-BmgBI | GCGGATCCGGAGATCTGACGGGTAATAACTGATATA | |
| NHM30 | TTGCCATTTATTGTCGCAGT | |
| JK149 | TCCCATAACCTCCTATATTGACTG | |
| NHM29 | CTTGGCCCTCTCCTTTTCTT | |
| NHM27 | CTGCTTCAAACCGCTAACAA | |
| JH53_Gib_mamminsert _F | CCCAATTCTGCAGATATCCATCACACTGGCGGCCGCTCGAGTGCAGACCTCAAATTCGAT | To amplify $(A_4G)_{60}$ from pJH4 to use Gibson assembly to replace pJH5 insert with $(A_4G)_{60}$ |
| JH54_Gib_mamminsert _R | TAGTAGGTTGAGGCCGTTGAGCACCGCCGCCGCAAGGAATGGTGCATGCTCGATGTGCAG | |
| JH55_Gib_mammvecto r_F | CTGCACATCGAGCATGCACCATTCCTTGCGGCGGCGGTGCTCAACGGCCTCAACCTACTA | To amplify vector from pJH5 to use Gibson assembly to replace pJH5 insert with $(A_4G)_{60}$ |
| JH56_Gib_mammvecto r_R | GCCAGTGTGATGGATATCTGCAGAATTGGGACCGTGCAATTCTTCTTACAGTTAAATGGG | |
| RM298 | TGCGGGTGTATACAGAATAGCAGAATGGGCAGACATTAC | |
| RM299 | CAATTCTAGATGCAGGCCGATCATCGTCGCGCTC | |

**Supplemental Table 6. Yeast strains used in this study**

| Strain | Genotype | Notes |
|---|---|---|
| JAH231 | *MATa leu2-Δ1, trp1-Δ63, ura3–52, his3–200, bar1Δ* | Background strain for yeast 2D |
| JAH315 | JAH231 2D plasmid JAH65 cl. 1A | |
| JAH316 | JAH231 2D plasmid JAH65 cl. 2A | |

| | |
|---|---|
| JAH317 | JAH231 2D plasmid JAH65 cl. 3B |
| JAH318 | JAH231 2D plasmid JAH69 cl. 1A |
| JAH319 | JAH231 2D plasmid JAH69 cl. 2A |
| JAH320 | JAH231 2D plasmid JAH69 cl. 3A |
| JAH321 | JAH231 2D plasmid JAH83 cl. 1A |
| JAH322 | JAH231 2D plasmid JAH83 cl. 2A |
| JAH323 | JAH231 2D plasmid JAH83 cl. 3A |
| JAH480 | JAH231 2D plasmid $(A_4G)_{60}$ cl. 2B |
| JAH481 | JAH231 2D plasmid $(A_4G)_{60}$ cl. 4B |
| JAH483 | JAH231 2D plasmid $(A_4G)_{60}$ cl. 17B |
